# Title: Time to wake up: Studying neurovascular coupling and brain-wide circuit function in the un-anesthetized animal

**DOI:** 10.1101/077024

**Authors:** Yu-Rong Gao, Yuncong Ma, Qingguang Zhang, Aaron T. Winder, Zhifeng Liang, Lilith Antinori, Patrick J. Drew, Nanyin Zhang

**Author notes:** Correspondence to: Patrick Drew, PhD Department of Engineering Science & Mechanics, Department of Neurosurgery W-317 Millennium Science Complex Pennsylvania State University University Park, PA 16802 or Nanyin Zhang, PhD Department of Biomedical Engineering W-341 Millennium Science Complex Pennsylvania State University University Park, PA 16802.

## Abstract

Functional magnetic resonance imaging (fMRI) has allowed the noninvasive study of task-based and resting-state brain dynamics in humans by inferring neural activity from blood-oxygenation-level dependent (BOLD) signal changes. An accurate interpretation of the hemodynamic changes that underlie fMRI signals depends on the understanding of the quantitative relationship between changes in neural activity and changes in cerebral blood flow, oxygenation and volume. While there has been extensive study of neurovascular coupling in anesthetized animal models, anesthesia causes large disruptions of brain metabolism, neural responsiveness and cardiovascular function. Here, we review work showing that neurovascular coupling and brain circuit function in the awake animal are profoundly different from those in the anesthetized state. We argue that the time is right to study neurovascular coupling and brain circuit function in the awake animal to bridge the physiological mechanisms that underlie animal and human neuroimaging signals, and to interpret them in light of underlying neural mechanisms. Lastly, we discuss recent experimental innovations that have enabled the study of neurovascular coupling and brain-wide circuit function in un-anesthetized and behaving animal models.

## Introduction

The discovery of anesthetics has been a boon to mankind, allowing invasive surgical procedures to be performed with little pain. In the history of neurophysiology, anesthetics were widely used to immobilize animals and to reduce variability in neural responses due to behavioral and attentional changes. Neurophysiological studies using the anesthetized preparation has tremendously advanced our understanding of brain function. Anesthetics are also an important tool for investigating the phenomenon of consciousness and theory of mind (Alkire et al., 2008; Brown et al., 2010; Brown et al., 2011), given their remarkable ability to manipulate consciousness level.

Despite the significant role that anesthesia plays in neuroscience research, it has become increasingly clear that anesthetics produced a neurological ‘state’ unlike any natural physiological condition, but rather, anesthesia is more akin to a ‘temporary, reversible coma’ (Brown, 2010). As a consequence of this better understanding of the nature of the anesthetized brain, and improvements in experimental methodology, there has been a push in the neuroscience community to utilize awake animal models in neurophysiological experiments (Ferenczi et al., 2016). Such models can be particularly valuable to the neurovascular coupling and animal neuroimaging communities, because the interpretation of both task-based and resting-state functional magnetic resonance imaging (fMRI) data collected in un-anesthetized humans depends on our understanding of neurovascular coupling under normal physiological conditions, and consequently, it is critical for translational animal experiments to be done without anesthesia as well. Furthermore, the vast majority of animal models of brain disorders rely on behavioral assessment conducted in the awake condition. Therefore, it is essential to image un-anesthetized animals if the imaging data will be used to interpret behavioral measurements (Ferenczi et al., 2016; Liang et al., 2014).

In this review, we summarize a large body of literature showing that anesthetics produce profound changes in cerebral hemodynamics, brain metabolism, neural activity, neurovascular coupling, and functional connectivity relative to the awake or sleeping states. Because of the large physiological differences between anesthetized and un-anesthetized animals, it is inappropriate to generalize physiological experimental data from anesthetized animals to physiological observations in awake humans. Given that technical advances have made it possible to study neurovascular coupling and brain circuit function in awake, behaving animals, the time is right for the community to move towards paradigms using awake animals.

### Physiological basis of the BOLD signal, and its sensitivity to anesthetics

The changes in oxygenation that underlie the blood-oxygenation-level dependent (BOLD) fMRI signal are due to an ‘oversupply’ of oxygenated blood to active areas of the brain (Buxton, 2012; Kim and Ogawa, 2012) (**Figure 1**). Neural activity directly and indirectly leads to the release of vasoactive substances (Attwell et al., 2010; Cauli and Hamel, 2010), which relax contractile tissue around arteries (Kim and Ogawa, 2012) and potentially capillaries (Hall et al., 2014; Hartmann et al., 2015; Stefanovic et al., 2008, Fernández-Klett et al., 2010; Hill et al., 2015). Because the resistance of a blood vessel depends on its diameter, these dilations cause a sharp reduction of vascular resistance, which increases blood velocity and flux. This increase in arterial diameter and blood flow is relatively rapid, occurring within less than a second following the stimulation (Chen et al., 2011; Drew et al., 2011; Gao et al., 2015; Kim et al., 2013). Arterial dilation and increased blood flow result in an elevated influx of oxygenated hemoglobin that exceeds the oxygen demand of the surrounding neural tissue (Fox and Raichle, 1986). Once the oxygenated blood transits through the capillary bed and into the draining veins, the increase in venous blood oxygenation can be detected using BOLD fMRI (Kim and Ogawa, 2012) (**Figure 1**). The BOLD signal depends on an interaction between local neural metabolic activity (which may not perfectly match up with the spike rate of neurons), increases in blood flow, volume and oxygenation. Because of the transit time, the BOLD signal is delayed relative to the increase in blood volume (Hirano et al., 2011; Silva et al., 2007a). A slow dilation of the veins underlies the “balloon” or “windkessel” models of the BOLD responses (Buxton et al., 2004; 1998; Mandeville et al., 1999; Miller et al., 2001). If the stimulus is sustained for tens of seconds, a slow dilation of veins is observed (Drew et al., 2011; Huo et al., 2015a; Kim and Kim, 2011; Mandeville et al., 1999), though the venous dilation seems to fail under some anesthetics (Drew et al., 2011; Hillman et al., 2007; Lee et al., 2001). Studying the relationship between neural activity and vascular dynamics (neurovascular coupling) is crucial for interpreting brain functional imaging signals. Substantial amount of previous work using anesthetized animals has significantly contributed to our understanding of neurovascular coupling and mechanisms of neuroimaging signals. However, as all of the processes that underlie the BOLD response are affected by anesthesia (**Figure 1**) (Aksenov et al., 2015; Martin et al., 2006b; Pisauro et al., 2013; Alkire et al., 2000; Lyons et al., 2016; Sellers et al., 2015; Ferezou et al., 2007; Nimmerjahn et al., 2009; Thrane et al., 2012; Chapin and Woodward, 1981; Crane et al., 1978; Cazakoff et al., 2014; Constantinople and Bruno, 2011; de Kock and Sakmann, 2009; Dudley et al., 1982; Ueki et al., 1992; Ferenczi et al., 2016), measurements made in the awake animal are expected to improve the quantification of neurovascular coupling and facilitate the translation over into the awake human studies.

**Figure. 1.**
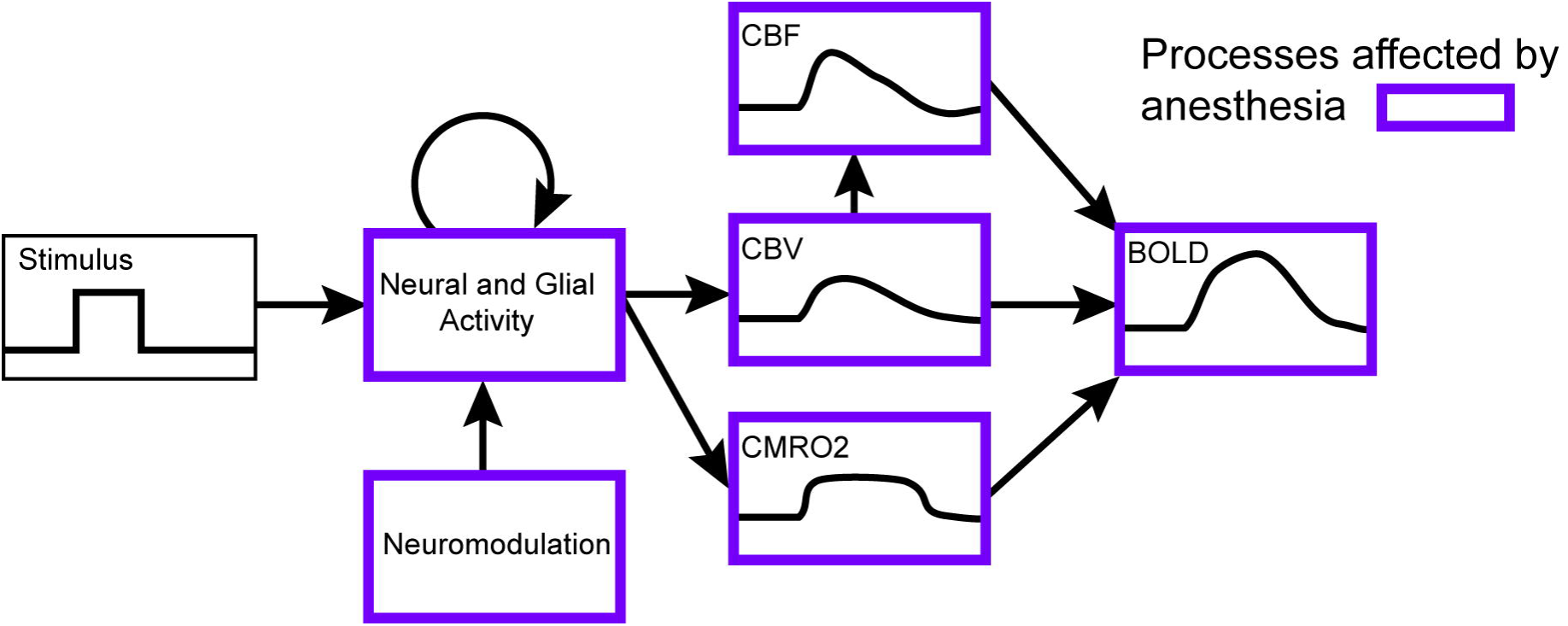
Effects of anesthetics on hemodynamic responses. Schematic showing how a stimulus-generated changes in neural activity are converted into a detectable BOLD signal. Physiological processes affected by anesthesia are outlined in purple.

### Anesthetics attenuate the amplitude and increase the lag of the hemodynamic responses

Hemodynamic signals are usually modeled as a convolution of local neural activity with a hemodynamic response function (HRF) to generate an observed blood volume or oxygenation change (Boynton et al., 1996; Logothetis et al., 2001; Vazquez and Noll, 1998). The amplitude and temporal dynamics of vascular dilations that follow increases in neural activity will determine the hemodynamic signals measured (Kim and Ogawa, 2012). Essentially all the processes that contribute to the formation of the HRF are suppressed by anesthesia (**Figure 1**). Importantly, though anesthesia reduces neural activity, it will disproportionally suppress the measured hemodynamic response (Logothetis et al., 2001; Goense and Logothetis, 2008; Pisauro et al., 2013; Aksenov et al., 2015). For instance, in the somatosensory cortex, fentanyl and/or isoflurane decrease multi-unit activity (MUA) and the BOLD response, however, the decrease in the BOLD response is substantially larger than the decrease in neural activity (Aksenov et al., 2015), implying a reduction in HRF amplitude. In addition, Pisauro et. al. directly measured MUA and cerebral blood volume (CBV) responses in awake and anesthetized mice (Pisauro et al., 2013), and found that the spike rates to visual stimulation in awake animals were slightly increased relative to anesthetized animals (~20%), but the hemodynamic response was increased to a much larger extent (~100%). This disproportional change again means that the amplitude of the HRF is reduced. A similar effect of anesthesia on the amplitude of the HRF is also seen both for cerebral blood flow (CBF) and CBV when neural activity is measured using the local field potential (LFP) (Martin et al., 2006b).

Anesthesia also causes a *slowing* in the HRF. Pisauro et al., observed a 2 second delay in the HRF in anesthetized mice relative to awake animals (Pisauro et al., 2013). This delay cannot be accounted for by a slight slowing in neural dynamics caused by anesthesia. The same group of researcher also reanalyzed the data of Logothetis et al. (2001) and Goense et al. (2008) and found a similar slowing of hemodynamic response in the anesthetized monkey relative to the awake monkey (Pisauro et al., 2013). Further, Martin and colleagues observed a similar slowing in the CBV, CBF and oxygenated hemoglobin responses in anesthetized rats relative to awake rats (Martin et al., 2006b). Taken together, these literature studies have found the HRF, the quantitative relationship between neural activity and blood flow, oxygenation or volume, is slowed in speed and decreased in amplitude by anesthesia (**Figure 2**).

**Figure. 2.**
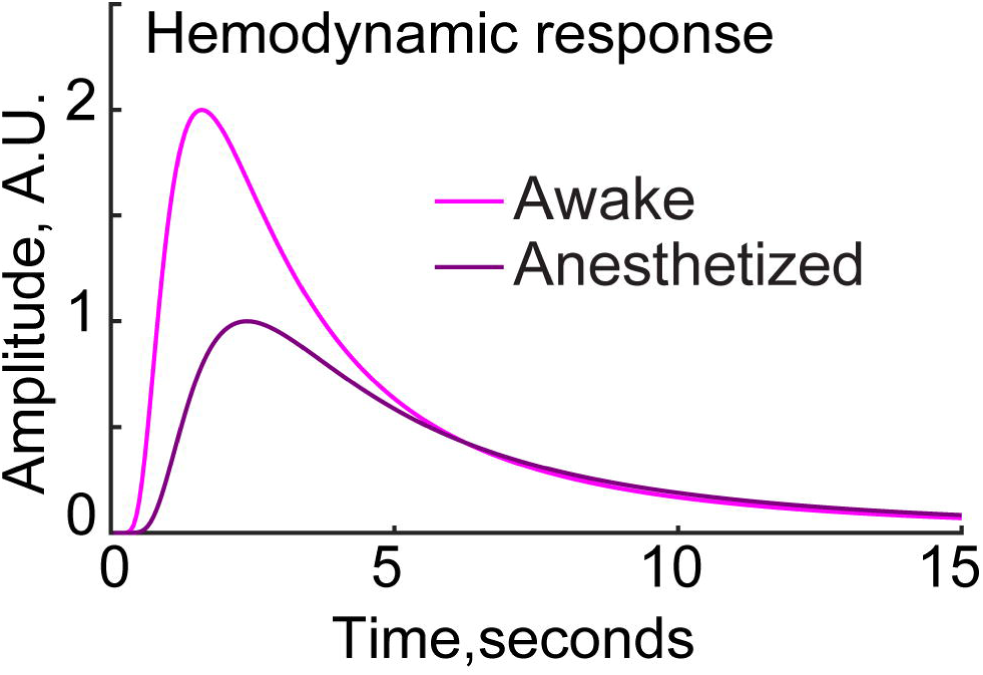
Schematic showing net effects of anesthesia on (BOLD) hemodynamic response function. Anesthesia slows and attenuates the hemodynamic response function. The awake hemodynamic response is approximately twice as large and has a faster onset compared to the HRF in anesthetized animals (Logothetis and Goense 2008; Pisauro et al., 2013; Aksnov et al., 2015).

Critically, an accurate, quantitative understanding of the relationship between neural activity and hemodynamic signals is critical for interpreting neuroimaging data. Handwerker and colleagues (2004) showed that slight variations in the time course of the hemodynamic response (1 second) can lead to erroneous fMRI statistical results. This difference is smaller than the 2 second delay difference found between awake and anesthetized animals (Pisauro et al., 2013). Therefore, to faithfully infer neuronal activity from hemodynamic signals, a quantitatively accurate HRF is essential, and anesthesia can represent a critical confounder in this aspect.

Current dominant working models for the vascular basis of neurovascular coupling (Buxton et al., 2004; 1998; Mandeville et al., 1999; Miller et al., 2001) lump the vasculature into ‘arterial’ and ‘post-arterial’ (capillaries and veins) components, assuming that changes in arterial volume are small, and venous distention dominate the bulk of CBV changes. However, using optical approaches, experiments with anesthetized animals had failed to detect any venous dilations (Hillman et al., 2007; Lee et al., 2001) and typically found small (~10%) average changes in arterial diameter in response to prolonged sensory stimulation (Devor et al., 2007; Nizar et al., 2013). In another two-photon imaging study, urethane blocked the dilation of veins (Drew et al., 2011). The lack of venous dilation, under some anesthetic regimens is likely due to the reduced blood pressure caused by anesthesia, as decreasing blood pressure in the awake animal blocks the venous dilation (Huo et al., 2015b). Interestingly, there are examples of putative slow venous dilations being detected using MRI (Kim and Kim, 2010; Silva et al., 2007a), though in both cases low doses of anesthetics were used.

Early optical imaging experiments revealed that in the awake animal, hemodynamic signals are substantially larger and more widespread in response to whisker stimulation (Martin et al., 2006a) or limb stimulation (Lahti et al., 1999; Peeters et al., 2001) than in anesthetized animals. Similarly, in the visual cortex, hemodynamic signals during wakefulness are approximately twice as large as those in the anesthetized animal (Goense and Logothetis, 2008; Liu et al., 2013a; Shtoyerman et al., 2000). Two-photon microscopy in awake, head-fixed mice has shown that arterial dilations in response to sensory stimulation are much larger (two-fold or more) than those in the anesthetized animal (**Figure 3**). Because the resistance of a vessel is inversely proportional to diameter to the fourth power, a small change in vessel diameter can lead to a large change in blood flow. Our data (**Figure 3**) showed a 20% dilation in the arteries in awake animals, will cause a 100% increase in flow, while in anesthetized animals an 8% dilation was observed, which will cause a 36% increase in flow. The dilation of arteries is accompanied by a slow distention of veins, which can build to levels of 10% or more for prolonged stimuli (Drew et al., 2011; Huo et al., 2015a; Gao and Drew, 2016). Voluntary locomotion drives nearly identical dynamics and amplitudes of arterial and venous dilations as those evoked by passive stimulation (Huo et al., 2015a), suggesting that this pattern of arterial and venous dilation is an invariant feature of cerebral hemodynamics found across different behavioral and cardiovascular states when the animal is awake.

**Figure 3.**
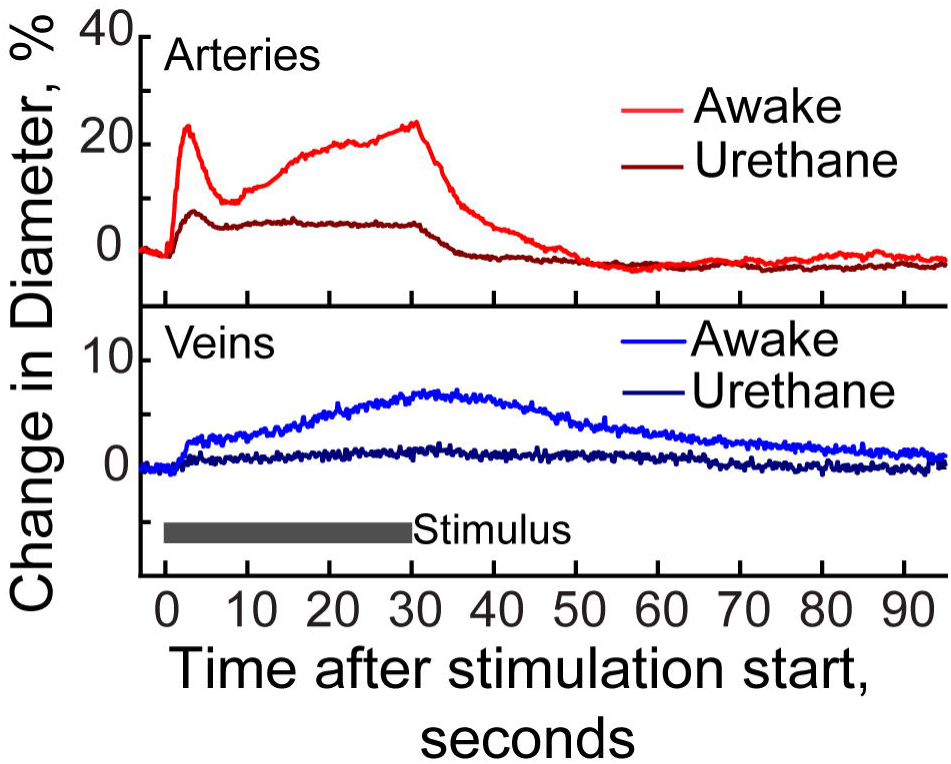
Comparison of dilation of individual arteries and veins measured using two-photon microscopy in the somatosensory cortex of awake and anesthetized mice. The sensory stimulus is puffs of air aimed at the vibrissae. Urethane drastically decreases the arterial dilation, and abolishes the venous dilation. Adapted from (Drew et al., 2011).

The anesthetic-induced reduction of hemodynamic response is at least in part caused by direct actions of anesthetics on ion channels of the vasculature. Like neurons, endothelial and smooth muscle cells have a rich compliment of ion channels, and many anesthetics have direct actions on these ion channels. The perhaps most well-studied anesthetic is the widely used vasodilatory anesthetic isoflurane. Isoflurane induces the relaxation of blood vessels via its action on ATP-sensitive potassium channels (Kokita et al., 1999), as well as reducing the calcium signals (Flynn et al., 1991) by blocking calcium channels (Buljubasic et al., 1992; Akata et al., 2003) in vascular smooth muscle. Some anesthetics, like halothane, cause vasodilation via an interaction with the endothelial cells to trigger the release relaxing factor (Muldoon et al., 1988), or via nitric oxide, like propofol (Park et al., 1995). These direct actions of anesthetics on ion channels present in the vasculature tend to have a shunting effect on the neurally induced currents, which can reduce the amplitude of the evoked dilation (Hill, 2012).

Lastly, the amplitude and dynamics of the hemodynamic response and its relationship to neural activity are greatly affected by the type and dosage of the anesthetic used (Masamoto and Kanno, 2012; Schlegel et al., 2015; Masamoto et al., 2009). For example, increasing isoflurane concentration (from 1% to 2%) does not measurably alter the neural responses to electrical simulation, but it does increase the stimulation-evoked blood flow response (Masamoto et al., 2009). Importantly, disproportionate influences of anesthesia on neuronal and vascular activities can lead to an altered neurovascular coupling relationship. Using optical imaging spectroscopy in awake rats, the neural–hemodynamic coupling relationship was found to be linear (Martin et al., 2006). This result is consistent with an approximately linear neurovascular coupling relationship found using fMRI in the microvasculature of awake humans (Zhang et al., 2008a; Zhang et al., 2009; Zhang et al., 2008b). However, this linear neurovascular coupling relationship was not present in anesthetized animals (Martin et al., 2006). Consequently, the results obtained under one anesthetic agent or dose are often not comparable to those obtained under another anesthetic agent or dose (Masamoto and Kanno, 2012; Schlegel et al., 2015; Masamoto et al., 2009). The issue as to which is the ‘correct’ anesthetic to use can be completely bypassed by working in the awake animal.

### Anesthetics cause profound decreases in baseline metabolism of the brain

The brain has an intense metabolic demand (Harris and Attwell, 2012), which is thought to constrain the computations neurons can perform (Attwell and Gibb, 2005; Howarth et al., 2012; Niven and Laughlin, 2008). Much of this energy is likely spent on recycling synaptic vesicles (Harris and Attwell, 2012; Rangaraju et al., 2014), with spiking activity and restoring the membrane gradient contributing less to energy consumption. Because glucose is the primary energy source for the brain (Siesjö, 1978), the most basic comparison across brain states is how the rate of glucose utilization (CMR_glc_) changes. CMR_glc_ can be measured in animal models using the 2-deoxyglucose (2-DG) technique (Sokoloff et al., 1977), or in humans using positron emission tomography (PET) (Reivich et al., 1979) or ^13^C MR spectroscopy techniques (Rothman et al., 2011).

Anesthetics cause a substantial depression of the resting metabolic rate in the brain (**Figure 4**), with the decreases in the metabolic rates in the cortex being larger than those in other brain structures. For example, alpha-chloralose, which is widely used in neurovascular coupling studies, causes a decreases of 50% or more in the metabolic rate of the cortex compared to the awake or lightly anesthetized state (Dudley et al., 1982; Hyder et al., 2002) with regional and layer variations in sensitivity (Dudley et al., 1982; Ueki et al., 1992). A dose dependent reduction of CMR_glc_ by pentobarbital is also observed, with a depression of 56% when cerebral pentobarbital levels exceed 50µg per g of tissue (Crane et al., 1978). In humans, halothane reduces global brain CMR_glc_ by ~40% (Alkire et al., 2000), and anesthesia with propofol (Alkire et al., 1995; Kaisti et al., 2003) or isoflurane (Alkire et al., 1997) cuts the cortical metabolic rate by 50% or more. To put this in context, one can compare anesthetic-induced decreases in brain metabolism to those observed in humans during sleep and those measured from patients in comas. Under normal physiological conditions, non-REM sleep decreases the brain metabolic rate by ~30% (Buchsbaum et al., 1989; Kennedy et al., 1982), while REM sleep decreases metabolic activity by ~10% (Buchsbaum et al., 1989), consistent with blood flow changes associated with sleep state (Braun et al., 1997). Patients in ‘locked-in’ and vegetative states showed decreases of 25% and >50% of brain metabolism rate, respectively (Levy et al., 1987). The large changes in basal metabolism will likely affect blood flow, as astrocytes, which play a key role in brain metabolism, exert control over tonic blood flow in the brain (Rosenegger et al., 2015).

**Figure 4.**
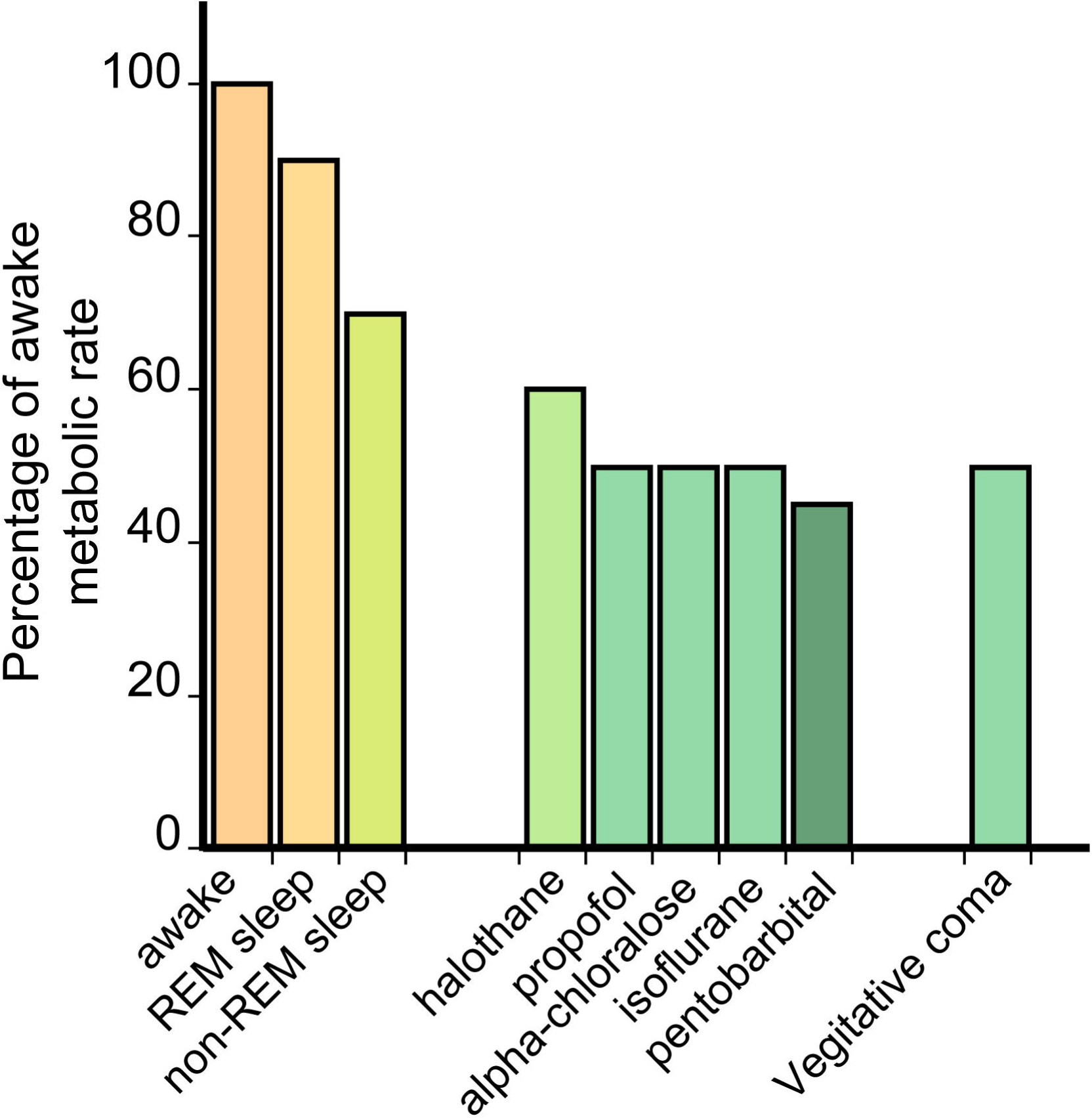
Brain metabolic rate is significantly decreased during anesthesia and sleep. Summary plot of measured brain metabolic rates under different anesthetics and conditions. References, species and measurement technique are as follows: REM sleep: Buchsbaum et al., 1989, Human 18-fluorodeoxyglucose PET. Non-REM sleep: Buchsbaum et al., 1989, Human 18-fluorodeoxyglucose PET, Kennedy et al., 1982, Rhesus monkey, 2-deoxyglucose. Halothane: Alkire et al., 1999, Human, 18-fluorodeoxyglucose PET. Propofol: Alkire et al., 1995, Human, 18-fluorodeoxyglucose PET. Propofol: Alkire et al., 1995, Human, 18-fluorodeoxyglucose PET. Alpha-chloralose: Dudley et al., 1982, Rat, 2-deoxyglucose. Isoflurane: Alkire et al., 1997, Humans, 18-fluorodeoxyglucose PET. Pentobarbotol: Crane et al., 1978, Rat, 2-deoxyglucose. Vegetative coma: Levy et al., 1987, Human, 18-fluorodeoxyglucose PET. Note that these are measures of changes in brain metabolism measured over the entire brain. Anesthesia-induced decreases in metabolic rate in gray matter will be larger than that in white matter because the metabolic rate of white matter is not appreciably affected by neural activity (Harris and Attwell, 2012).

Anesthesia-related brain metabolism changes can also affect oxygenation. Microscopic oxygen measurements using polarographic electrodes have shown great heterogeneity in tissue oxygen concentration in anesthetized animals (Ndubuizo and LaManna, 2007), making direct comparisons between the awake and anesthetized state difficult. Recent technical developments combining phosphorescent oxygen-sensitive dyes with two-photon microscopy have enabled the visualization of oxygen gradients at the micrometer scale (Sakadzic et al., 2010, 2014), showing that tissue oxygen tension varies widely depending on the distance to the nearest arteriole. Two-photon phosphorescence point measurements of oxygenation have found that the oxygen tension in the brains of awake mice are lower than in anesthetized mice, and have also failed to find laminar differences in oxygen levels which are seen in anesthetized animals (Lyons et al., 2016). Though further studies are warranted, these results suggest that the metabolic changes caused by anesthesia could alter normal brain oxygenation, which will have implications on BOLD signals measured in anesthetized animals.

### Anesthetics directly alter the responsiveness and dynamics of neurons and glia

There is a long history of using anesthetized animal models in sensory neurophysiology studies (Hubel and Wiesel, 1959), allowing the state of the animal to be controlled and the presentation of stimuli unaffected by eye movements, attention and other sources of variability present in awake animals. Gradually, it became apparent that neural dynamics during anesthesia were very different from those in the awake animal in several important ways. First, anesthetics, in general, tend to hyperpolarize neurons by increasing inhibition and/or decreasing excitation (Alkire et al., 2008; Ries and Puil, 1999), which usually leads to a suppression in the average firing rate of neurons (Cazakoff et al., 2014; Constantinople and Bruno, 2011; de Kock and Sakmann, 2009). Electroencephalgraphy (EEG) recordings show that baseline neuronal activity is suppressed by various anesthetics (Antunes et al., 2003; Austin et al., 2005; Bonvento et al., 1994), while the evoked neural response is different in shape and is significantly delayed under anesthesia (Martin et al., 2006b; Sellers et al., 2015). Higher order aspects of neural activity are also profoundly affected, as correlations between neurons are increased during anesthesia (Greenberg et al., 2008).

The receptive field of neurons in sensory areas is also altered by anesthesia. A common but not universal finding is that the receptive field of a neuron is reduced by anesthesia (Armstrong-James and George, 1988; Chapin and Woodward, 1981; Friedberg et al., 1999). This distortion will have implications for neuroimaging-based mapping experiments since the activated area following sensory stimulation may be much larger in the awake animal (Ferezou et al., 2007). However, in the visual cortex, anesthesia seems to increase the spatial spread and prolong the duration of the response to visual input (Haider et al., 2013; Vaiceliunaite et al., 2013). In the olfactory bulb, odor evoked responses are larger in the anesthetized animal (Kato et al., 2013; Rinberg et al., 2006). In higher order sensory and sensorimotor areas, anesthesia can have large effects on sensory gating and responsiveness (Cardin and Schmidt, 2004, 2003; Schmidt and Konishi, 1998). These complicated, and brain-region specific, changes make translation of imaging results obtained under anesthesia to the awake brain difficult.

Although anesthesia affects neural activities in many ways, these effects are not uniform across the brain, or even within the cortex. Frontal areas tend to be more affected by anesthesia than sensory areas (Sellers et al., 2013; Liang et al., 2015a), and there are laminar differences as well (Raz et al., 2014; Sellers et al., 2013). Anesthesia is also not a temporally uniform state. Many anesthetics, including urethane and isoflurane, are known to make the cortex alternate between a depolarized up-state and a hyperpolarized down-state (Castro-Alamancos, 2009; Steriade et al., 2001). In the ‘up’ state, neurons are more responsive to external input (Curto et al., 2009; Shu et al., 2003), similar to the sustained depolarization of wakefulness, while the synaptic activity is reduced during the ‘down’ state (Castro-Alamancos, 2009). Some effects of anesthesia may be due to changes in the modulatory state of the brain, as stimulation of modulatory nuclei can induce ‘awake like’ behavior. Stimulation of the nucleus basalis, which send cholinergic projections to cortex, can facilitate motor behavior (Berg et al., 2005) and alter sensory responses in anesthetized animals to a more ‘awake’ like state (Castro-Alamancos and Gulati, 2014; Goard and Dan, 2009). Anesthesia also increases the cortical adaptation to repetitive sensory stimulation relative to the awake state (Castro-Alamancos, 2004; Masamoto et al., 2009).

In addition to the depressant effects of anesthesia on neural activity, anesthetics decrease the frequency of calcium signals in astrocytes. Astrocytes are essential for the supply of energy metabolites to neurons, and provide a physical link to the vasculature (Simard et al., 2003; Takano et al., 2006). Astrocytes are also thought to have a role in mediating vasodilation in response to increased neural activity (Hirase, 2005; Takano et al., 2006), although this issue is contentious (see (Bonder and McCarthy, 2014; Lind et al., 2013; Nizar et al., 2013; Otsu et al., 2014)). Using two-photon microscopy, clear decreases in the spontaneous calcium signals in astrocytes have been observed under isoflurane, ketamine and urethane (Nimmerjahn et al., 2009; Thrane et al., 2012). In both studies, the anesthetic-induced astrocyte activity decrease was larger than the decrease induced by local pharmacological blocker of sodium channels or glutamate receptors, implying that the anesthetics have direct action on astrocytes. These effects of anesthesia on neural and glial activity are further complicated by the possibility that anesthesia could disrupt processes that relate neural activity changes to the hemodynamic response. For example, a study investigating the role of neuronally-generated nitric oxide (NO) in neurovascular coupling found different effects of NO synthase inhibitor on CBV in awake and anesthetized animals (Nakao et al., 2001), suggesting differential involvements of neurovascular coupling pathways depending on the state of the animal.

Lastly, due to the decreased responsiveness of neurons during anesthesia, many groups use electric shocks, rather than tactile or visual stimuli, to elicit hemodynamic responses. Electric shock is an unpleasant experience that bears little resemblance to normal sensory experience that indiscriminately activate proprioceptive, pain and other fibers, and can also induce large systemic changes in blood pressure that obscure neuronally-generated hemodynamic signals (Schroeter et al., 2014). As much larger (two-fold or greater) hemodynamic responses can be elicited in the awake animal (Drew et al., 2011; Pisauro et al., 2013) using stimuli that are more of the type that might be presented to a human during an fMRI experiment, the advantages of the awake preparation are clear.

### Anesthetics decrease brain temperature and affect other brain physiological processes

While neurophysiologist focus on electrical measures of neural activity, there are other physiological variables that are greatly perturbed by anesthesia that could potentially influence the neurovascular coupling. Anesthetics decrease brain temperature by several degrees (Hayward, 1968; Shirey et al., 2015; Kalmbach and Waters, 2012). The decrease in brain temperature persists even when the core body temperature is kept in the physiological range (Shirey et al., 2015; Kalmbach and Waters, 2012; Podgorski and Ranganathan, 2016). The decrease in brain temperature is largest at the surface of the brain (Kalambach and Waters, 2012; Podgorski and Ranganathan, 2016), potentially altering the dynamics of the large pial vessels and supragranular neurons more than the intraparenchymal vessels. Since neural activity warms the brain (Kiyatkin et al, 2002), the brain cooling associated with anesthesia is partially due to the anesthesia-evoked decreases in brain metabolism. Alternations of temperature, either up or down, affect the baseline diameter, myogenic tone, and spontaneous vasomotion of an arteriole (Osol and Halpern 1988; Ogura et al., 1991), which will change evoked vascular dynamics under anesthesia (Pisauro et al., 2013). Brain temperature changes of one degree or more are large enough to produce changes in animal behavior (Long and Fee, 2008; Aronov and Fee 2012).

Another physiological change induced by anesthesia is an increase in the volume fraction of extracellular space. The brain is composed of approximately 20% extracellular space (Nicholson, 2001; Korogod et al., 2015). The extracellular volume fraction has been shown to increase relative to the awake condition under ketamine (Xie et al., 2013). Because the extracellular space exchanges ions with active neurons, changes in the volume of this extracellular space should impact neural excitability (Somjen, 2002) and cerebral spinal fluid flow (Iliff et al., 2013). Besides brain temperature and extracellular space, intracranial pressure (ICP) is another important physiological variable that can be altered by various anesthetics (Artru, 1984; Gao and Drew, 2016). Elevating ICP alters heart beat and respiratory rhythm (Matsuura et al., 1984), decreases CBF in the cortex (Sadoshima et al., 1981), and changes neural activity (Seigo Nagao et al., 2009), all of which could impact neurovascular coupling. Lastly, isoflurane can open the blood-brain barrier, leading to ionic changes in the brain (Chever et al., 2008). While these physiological changes by anesthesia may not be as commonly appreciated as neural or cardiovascular changes, they are likely very disruptive.

### Anesthesia alters neural circuit function

Investigating neural circuit function plays a fundamental role in neuroscience. The brain is a highly inter-connected system that gives rise to complex, diverse and integrated information processing (Alkire et al., 2008; Tononi et al., 1998). The proper function of neural circuitries and networks is essential to maintain meaningful information exchange and integration, and thus to support normal brain function. Elucidating the functional properties of neural networks, and their dysfunction in the context of pathology, is a critically important avenue of investigation in neuroscience.

Neural circuit function can be studied at both resting and activated states using fMRI techniques. In the resting state, fMRI (termed resting-state fMRI or rsfMRI) measures resting-state functional connectivity (RSFC) between brain regions by quantifying the temporal synchronization of spontaneously fluctuating rsfMRI signals (Biswal et al., 1995; Fox and Raichle, 2007). This technique offers a convenient tool to study neural circuit function without relying on any external stimulation, and thus is particularly suitable in some clinical settings where patients have difficulty to perform certain tasks inside the MRI scanner. Indeed, RSFC in awake humans has been extensively used to reveal neural circuit function in both normal (Albert et al., 2009; Horovitz et al., 2009) and pathological conditions (Anand et al., 2005; Carter et al., 2012; Greicius et al., 2007; Greicius et al., 2004; Hunter et al., 2012; Kennedy et al., 2006; Lowe et al., 2002; Lustig et al., 2003; Mayer et al., 2011; Tian et al., 2006; van Meer et al., 2012; Whitfield-Gabrieli et al., 2009).

Anesthesia has been shown to profoundly alter RSFC measurements of neural circuit function. A number of animal studies indicate that RSFC is tremendously weakened by anesthesia. Lu *et. al.* demonstrated a dose-dependent decrease of cross-hemispheric RSFC in alpha-chloralose-anesthetized rats (Lu et al., 2007). This finding is in agreement with the study by Liu and colleagues who found that intrinsic BOLD fluctuations and functional connectivity in the resting rat are strongly dependent on anesthesia depth (Liu et al., 2011). In addition, several studies have shown that, relative to the awake state, thalamo-cortical connectivity is significantly reduced by anesthesia in the rat (Liang et al., 2012b; Liang et al., 2013). This result is consistent with extensive literature demonstrating a pivotal role of the thalamus in the systems-level mechanisms of anesthetic-induced unconsciousness (Nallasamy and Tsao, 2011). Interestingly, there is evidence indicating that thalamo-cortical connectivity in higher-tier associative networks is more strongly affected than that in lower-level sensory-motor networks by anesthesia, highlighting the importance of consciousness on integrative multimodal processing (Boveroux et al., 2010; Liang et al., 2012b; Liang et al., 2013). Furthermore, dynamic rsfMRI approaches reveal that anesthesia diminishes RSFC between medial prefrontal cortex (mPFC) and the hippocampus (Liang et al., 2015a), as well as RSFC between mPFC and amygdala (Liang et al., 2012b) as compared to the awake state, suggesting that anesthesia can profoundly impact neural circuits subserving cognitive and emotional function. This result is consistent with the notion that cognitive and emotional regulations are overall disrupted in the anesthetized state.

Similar effects of anesthesia on RSFC have also been observed in numerous human studies. RSFC in the cortex is ablated in deeply anesthetized patients (Peltier et al., 2005). In particular, like animal studies, anesthesia disrupts RSFC within the thalamocortical and frontoparietal networks in humans (Breshears et al., 2010; Lee et al., 2009; Velly et al., 2007; White and Alkire, 2003). Like anesthesia, sleep has similar effects on RSFC (Tomasi et al., 2009). Spontaneous fluctuations of the BOLD signal and RSFC are significantly reduced in sleep compared to wakefulness (Wilson et al., 2015).

Anesthesia can also lead to considerable changes in anticorrelated RSFC. Although the majority of rsfMRI studies use positive temporal correlations between BOLD time series as a measure of RSFC, negative correlations (i.e. anticorrelated RSFC) also contains important information of neural circuit function (e.g. inhibitory interactions between regions). For instance, anticorrelated RSFC has been shown to play a critical role in mediating cognitive function in humans (Fox et al., 2005; Hampson et al., 2010; Kelly et al., 2008). In animal studies, robust anticorrelated RSFC in a well-characterized frontolimbic circuit between the mPFC and amygdala exhibited in the awake rat, whereas this anticorrelated RSFC was abolished in anesthetized rats (Liang et al., 2012a) (**Figure 5**). These results are consistent with a recent study reporting both positive and negative RSFC in the awake monkey, but only positive RSFC when the monkey was anesthetized (Barttfeld et al., 2015).

**Figure 5.**
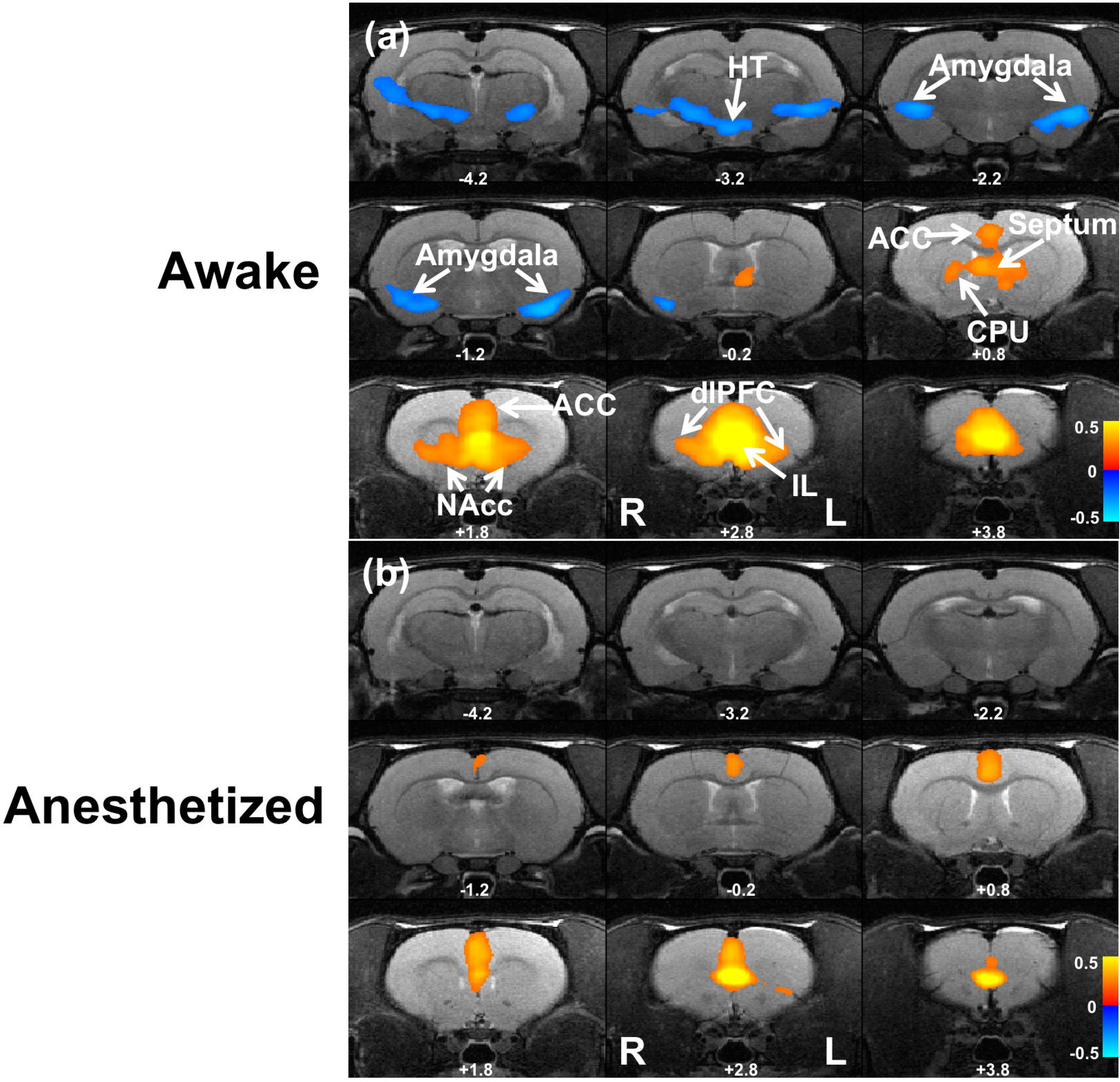
Infralimbic cortex (IL) resting-state functional connectivity maps in awake and anesthetized rats. (a) The IL RSFC map in the awake rat. (b) The IL RSFC map in the anesthetized rat. Anticorrelated RSFC between IL and amygdala was completely abolished by anesthesia. All maps were overlaid on anatomical images. Distances to Bregma (mm) are labeled for each slice. Adapted from (Liang et al., 2012a).

In addition to the resting state, neural circuit function can be studied in activated states. The emergence of optogenetics has opened a new avenue for interrogating the neural circuit function that mediates many behaviors (Deisseroth et al., 2006). Such capacity has been highlighted by a series of recent studies that combine optogenetic control and fMRI readout (Abe et al., 2012; Kahn et al., 2013; Lee et al., 2010; Li et al., 2011; Liang et al., 2015b; Weitz et al., 2015). In an optogenetic approach, light-activated ion channels are introduced to a specifically targeted neuron population through adeno-associated viral (AAV) injection. This allows researchers to specifically activate/inhibit the infected neurons through optical stimulation (Deisseroth et al., 2006). The scientific value of this technique is tremendously expanded by combining it with fMRI (opto-fMRI), because the signaling impact of photo-activating a specific circuit element on the rest of circuit can be monitored across the whole brain (Lee et al., 2010). Compared to conventional electrical stimulation approaches, opto-fMRI does not suffer from electromagnetic-related artifacts that electrical approaches commonly encounter.

Anesthesia also exerts significant effects on neural circuit function in the activated state. Opto-fMRI conducted in awake mice (Desai et al., 2011) and rats (Ferenczi et al., 2016; Liang et al., 2015b) have consistently indicated that BOLD activations accompanying optogenetic stimulation in both injected regions and postsynaptic targets are strongly suppressed by anesthesia. In addition, the neural circuit connectivity pattern is significantly altered in the anesthetized state (Liang et al., 2015b) (**Figure 6**). Taken together, all these studies strongly suggest that anesthesia can significantly alter neural circuit function. In fact, RSFC changes in anesthetized states have been detected using rsfMRI and have provided important insight into the system-level neural mechanism underlying anesthetic-induced unconsciousness (Liang et al., 2012b).

**Figure 6.**
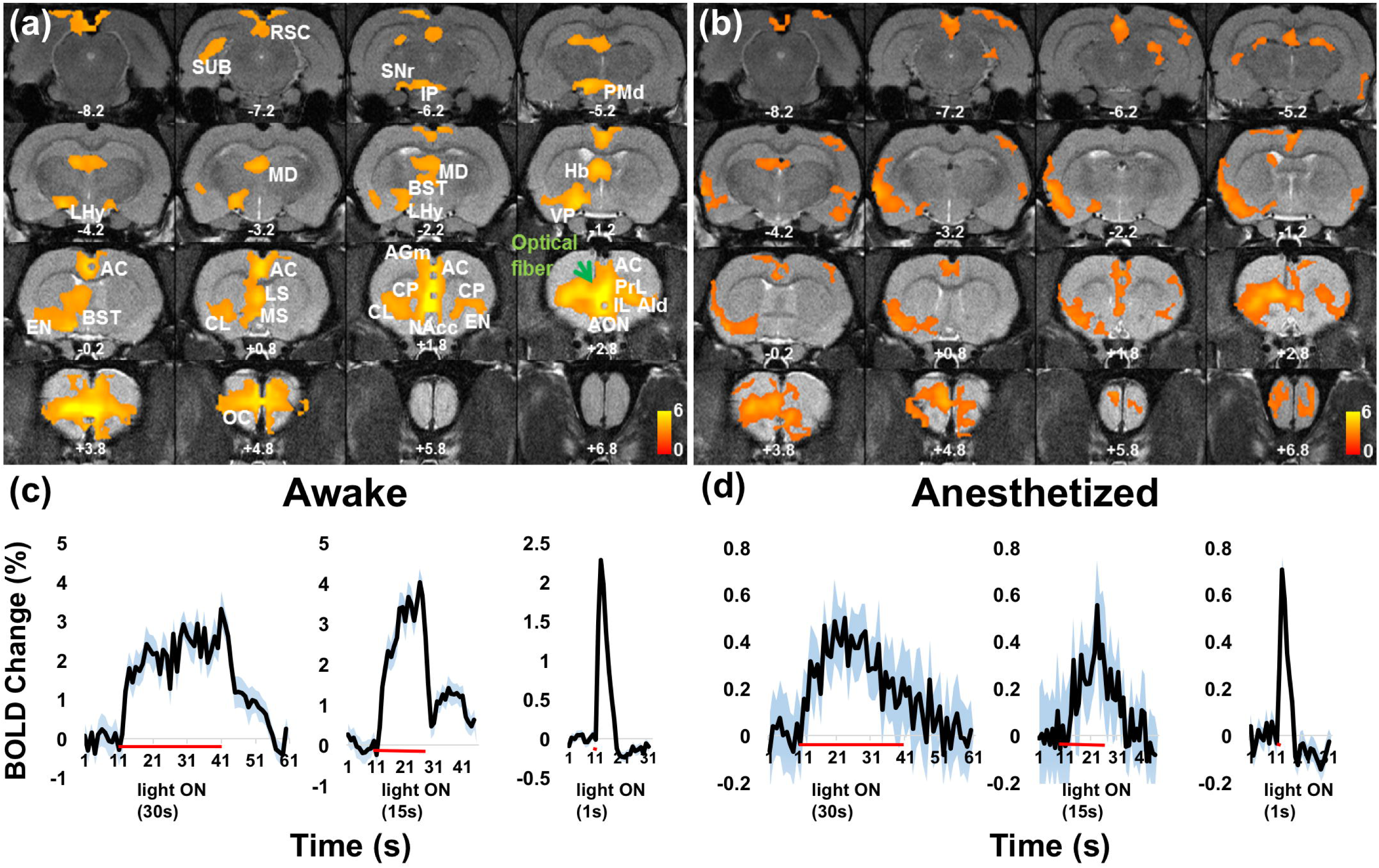
Optogenetic activation of brain networks in the awake and anesthetized animal. Averaged activation maps and time courses in response to infralimbic (IL) optogenetic stimulation in awake (left) and isoflurane-anesthetized (right) states. a) the averaged activation map in the awake state. b) the averaged activation map in the anesthetized state. Green arrow, implanted optical fiber. Distance to Bregma is labeled in each slice. c) BOLD time courses of anterior cingulate cortex in the awake state. d) BOLD time courses of anterior cingulate cortex in the anesthetized state. Note the difference in the vertical scale. Red lines indicate the stimulation periods. Blue shades shows SEM. AC, anterior cingulate cortex; AGm, medial agranular (frontal) cortex; AId, dorsal agranular insular cortex; AON, anterior olfactory nucleus; BST, Bed nucleus of stria terminalis; CL, claustrum; EN, endopiriform nucleus; CP, caudate-putamen; Hb, habenula; IL, infralimbic; IP, interpeduncular nucleus; LHy, lateral hypothalamus; LS, lateral septum; MD, mediodorsal nucleus; MS, medial septum; NAcc, nucleus accumbens; OC, orbital cortex; PMd, dorsal premammillary nucleus; PrL, prelimbic; SNr, substantia nigra; SUB, subiculum; RSC, retrosplenial cortex. Adapted from (Liang et al., 2015b).

### Anesthesia causes changes in brain network organization

The intrinsic functional connectivity architecture is a crucial component of the governing principles of brain organization. This connectivity architecture has typically been studied by combining rsfMRI (or diffusion tensor imaging) with *graph theory*. Brain graphs are composed of a number of *nodes* interconnected by a set of *edges*. Anatomically or functionally defined region of interests (ROIs) are defined as nodes in the brain graph, and functional or structural connectivity between ROIs is used to define edges. Connectional architecture of brain networks can be quantitatively evaluated using a set of topological parameters of the brain graph, such as *degree, clustering coefficient, shortest path length and modularity*.

Relative to the awake state, brain networks are topologically altered to support different patterns of information transfer in the anesthetized state. Our data indicate that neural networks are significantly reorganized and exhibit different network properties (Liang et al., 2012b). For instance, several brain regions demonstrate pronounced changes in local topological metrics such as *clustering coefficient* and *betweeness centrality*. In particular, relative to the awake condition, regions of the basal ganglia, including nucleus accumbens and septal nuclei, show significantly reduced local clustering coefficients in the isoflurane-induced anesthetized condition, which suggests that the communication between those regions and their neighboring regions is reduced in the anesthetized animal. Interestingly, the same two regions were reported to enhance anesthetic effects when they were pharmacologically inactivated (Ma and Leung, 2006; Ma et al., 2002). These results collectively highlight the susceptibility of the basal ganglia to anesthesia. In addition, thalamic nuclei show decreased betweenness centrality, again indicating impaired information relay in the thalamus in the anesthetized brain. Furthermore, the community structure between the awake and anesthetized states considerably differs in ways with stark functional implications (**Figure 7**). Brain modules in the awake brain are more likely to contain both cortical and sub-cortical regions (Liang et al., 2011), whereas brain modules in the anesthetized brain tend to include only cortical or only subcortical regions, implying compromised communication between the cortex and subcortex in the anesthetized state (Liang et al., 2012b). The impact of anesthesia on the topological organization of brain networks has also been reported in several EEG studies. Lee et al demonstrated reconfiguration of network hub structure after propofol-induced unconsciousness (Lee et al., 2013), and a disruption of network topology during the transition to loss of consciousness (Lee et al., 2011). Taken together, these studies strongly indicate that brain networks are topologically reorganized to support new patterns of information integration in the anesthetized condition.

**Figure 7.**
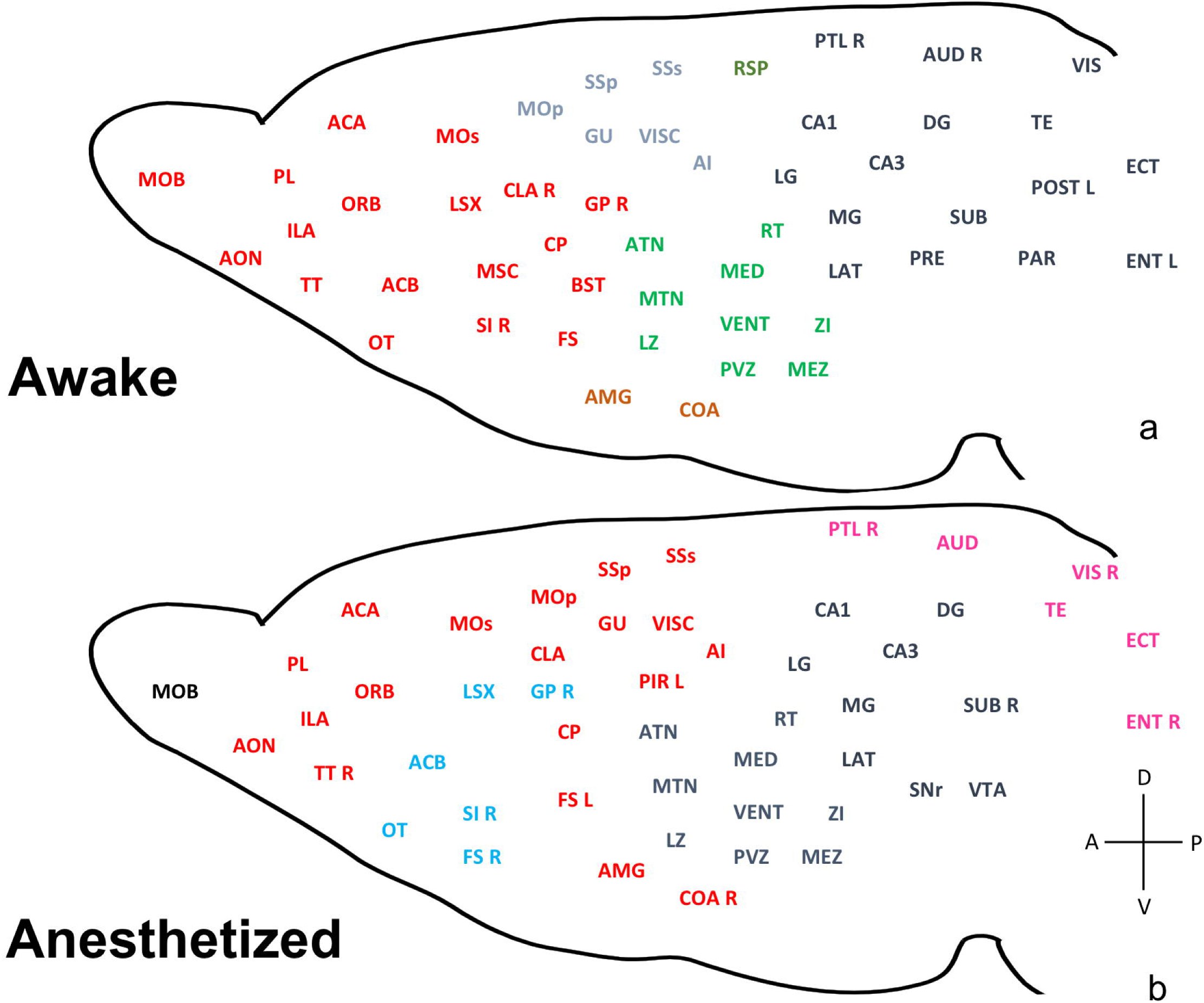
Anesthesia disrupts large-scale brain organization. Community structures in the awake (a) and isoflurane-anesthetized (b) states. Brain regions with the same color are within the same module. Brain regions without the annotation of L or R imply bilateral sides. L: left, R: right, A: anterior, P: posterior, V: ventral, D: dorsal. Adapted from (Liang et al., 2012b).

### Technical advances enable the study of neurovascular coupling and neural circuit function in the awake animal

In the systems neuroscience community, the realization that anesthesia profoundly disrupts normal neural circuit function has driven neurophysiologists to embrace the use of awake animals, and many of these approaches and tools can be translated to the study of neurovascular coupling. A key advance has been the widespread adoption of head-fixation methods, first developed in the primate neurophysiology community (Wurtz, 1969) and then adapted the use in rodents (Dombeck et al., 2007)(see also Kleinfeld and Griesbeck, 2005 and references therein). Both mice (Murphy et al., 2016) and rats (Scott et al., 2013; Scott et al., 2015) can be trained to voluntarily engage in head fixation, indicating that head fixation is not an aversive experience. Though hemodynamic signals can be imaged from awake, head-fixed rodents using fMRI or optical imaging, there are methodological differences due to technical considerations. Additionally, there are species differences, with mice acclimating to head-fixation more rapidly than rats, which in turn acclimate more rapidly than marmosets (Belcher et al., 2013b; Liu et al., 2013a) and other primates. Each species has its own unique advantages and limitations, and the choice of species will be driven by the scientific questions addressed (Hall et al., 2014; Hill et al., 2015; Hung et al., 2015a). Below we discuss the approaches used in mice and rats.

### Habituation to head fixation

The habituation procedures depend on species (mice or rats) and imaging modality (MRI or optical). Optical imaging requires a very still head relative to the optical imaging apparatus, requiring a metal headbar, but this type of fixation is avoided in MRI due to the issue of devices’ MRI compatibility. Anecdotally, mice are easier to acclimate to head fixation, with some groups being able to bypass acclimation all together (Dombeck et al., 2009; Sofroniew et al., 2014).

For optical imaging experiments, prior to habituation a headbar is surgically attached to the skull to hold the head still. A cranial window may also be implanted at this time. After allowing adequate time for surgical recovery, we begin the habituation to head-fixation, which can be done on treadmill that allows locomotion (Dombeck et al., 2010; Gao et al., 2015; Gao and Drew, 2014; Huo et al., 2014; Rosenegger et al., 2015; Tran and Gordon 2015), or body restraining cylinders or slings (Berwick et al., 2002; Drew et al., 2011) (**Figure 8**). We typically habituate mice to being on the treadmill or head-fixing in a tube by acclimating them over two to five days (Drew et al., 2010a; Drew et al., 2011; Huo et al., 2015a;Gao and Drew, 2016) in several 15-minute to ½-hour sessions. Mice are briefly (<1 min) anesthetized with isoflurane before being head fixed to minimize handling stress. This habituation procedure and schedule is based on those used in neurophysiology experiments (Dombeck et al., 2007; Zagha et al., 2013). With a stable preparation, the noise in the intrinsic signal due to animal motion is negligible, as it is several orders of magnitude below the amplitude of the sensory-evoked response (Huo et al., 2014; 2015a; 2015b). Two-photon microscopy is sensitive to animal movement, but movement during voluntary locomotion are typically small enough to be corrected using motion correction software (Dombeck et al., 2007; Gao and Drew, 2016). Head-fixation and optical imaging can be longitudinally done on very young animals, starting as early at postnatal day 3 (P3) (Letourneur et al., 2014). Using two-photon microscopy, we are able to image locomotion-induced vasodilation in the somatosensory cortex (**Figure 9**). Note that even small movements can drive 5-10% increases in arterial diameter, and prolonged motion can lead to arterial dilation of 40% or more, much larger than what is observed in the anesthetized animal. Large windows allow chronic measurement of hemodynamic signals over the entire dorsal surface of the cortex. The arterial dilation is localized to the somatosensory and visual cortex (**Figure 10**), with little arterial dilation seen in the frontal cortex (Huo et al, 2014, 2015a, 2015b, Gao and Drew, 2016).

**Figure 8.**
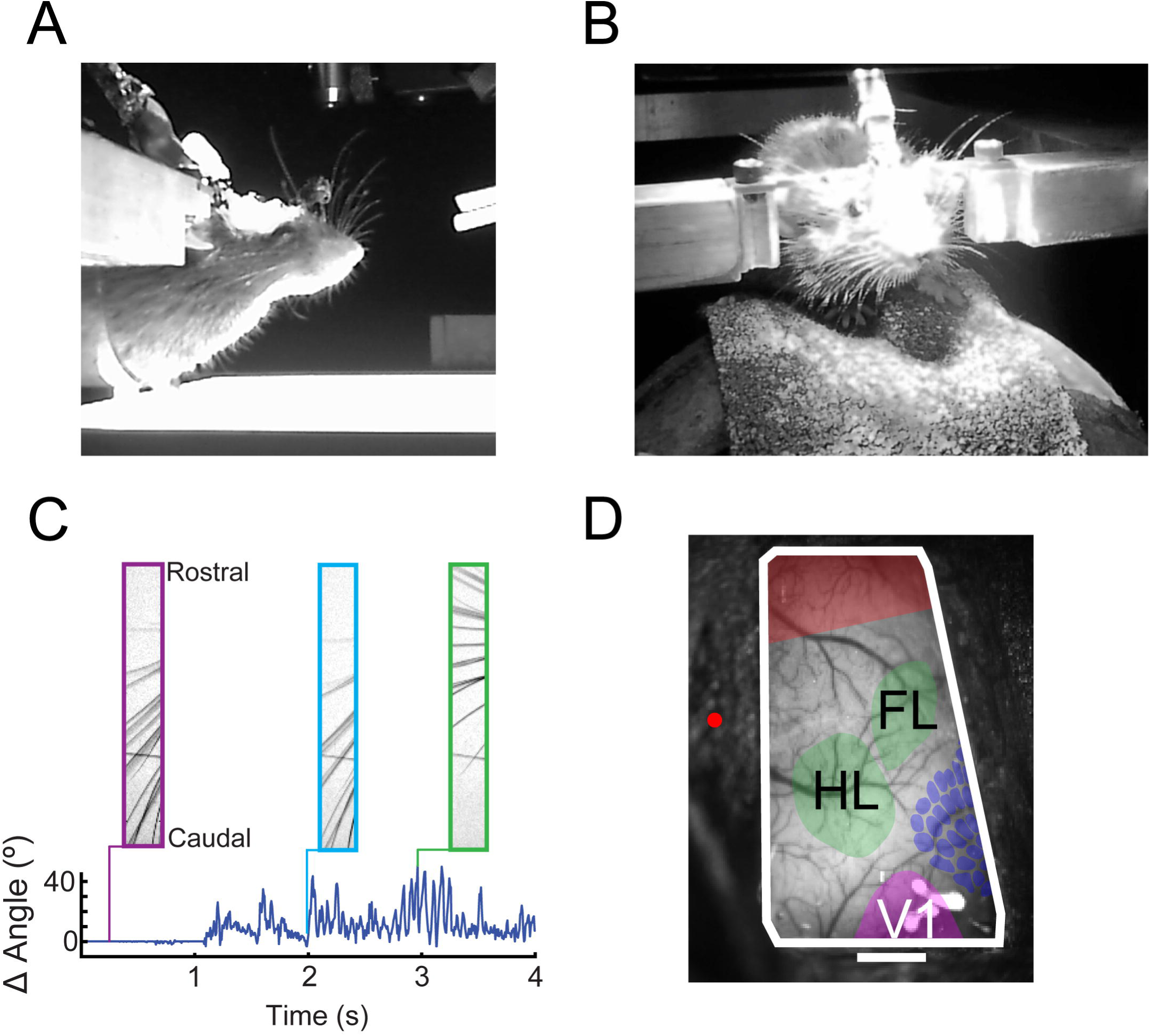
Imaging hemodynamic signals in awake mice. A) Mouse head-fixed in a plexiglass cylinder. The whiskers are illuminated from below to allow tracking of movements. B) Mouse head-fixed on a spherical treadmill with one degree of rotational freedom. The mouse is free to run or stand still. An electrical connector allows measurement of simultaneous ECoG or LFP from implanted electrodes. C) Head fixed mice exhibit active sensing (whisking) behavior. Top, camera images taken of whiskers of a head-fixed mouse at various time points and protraction angles. Bottom, whisker angle versus time tracked using the Radon transform (Drew et al., 2010b; Gao and Drew, 2014). D) The entire dorsal cortex can be imaged using optical imaging. Photo of a large, thinned skull window showing locations of forelimb/hindlimb representations (green, FL/HL), frontal cortex (red), visual cortex (purple, V1), and whisker representation (dark blue) reconstructed from the cytochrome oxidase stained cortex. Red dot denotes the location of bregma. Rostral is up, medial to the left. Scale bar 1 mm.

**Figure 9.**
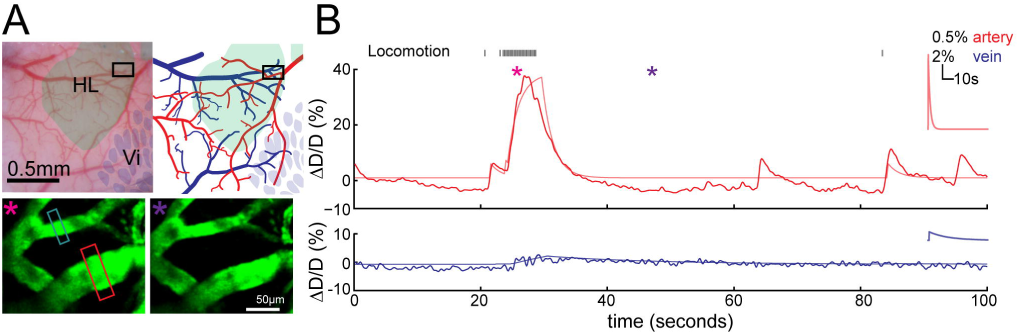
Cerebral arterial and venous responses during voluntary locomotion in head-fixed awake mice measured with two-photon microscopy. A) Top left, photograph of the cortical vasculature taken through a Polished and reinforced thinned skull (PoRTS) window. The window was implanted over the parietal cortex, which includes the hindlimb region (HL) and vibrissa cortex (Vi). The black box denotes the region where the two-photon images were obtained. Top right, schematic drawing of the major surface vasculatures on the PoRTS window in A. Arterioles are in red, while venules are in blue. Bottom, one second averaged two photon images of an arteriole (in red box) and a venule (in blue box) enclosed in the black box in A during resting (right, marked by purple asterisk) and locomoting (left, marked by magenta asterisk). The arteriole dilates strongly during locomotion. **B)** The responses of the arteriole (top) and venule (bottom) shown in C during voluntary locomotion. Black tick marks on top are the binarized locomotion events. Locomoting and resting periods where the two-photon images shown in A are marked by magenta and purple asterisks, respectively. Arteriole dilation is rapid and large, while venule diameter changes are smaller and slower. The dilations of both vessels can be captured by convolving the binarized locomotion events with an exponential decay (Gao and Drew, 2016; Huo et al., 2015a, 2015b). The fitted responses are shown in light red and light blue for arteriole and venule respectively. Insets are the fitted exponential kernels.

**Figure 10.**
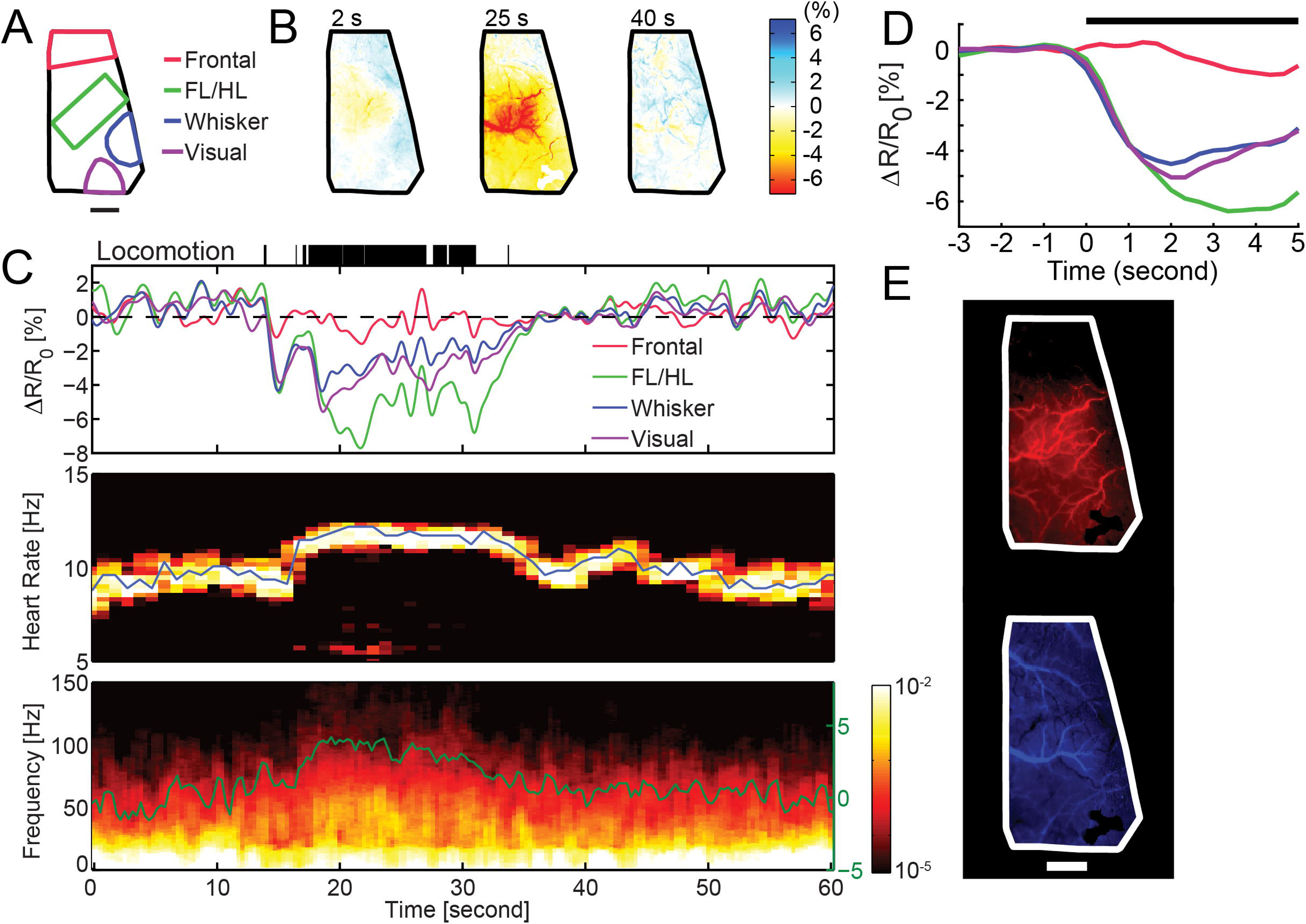
Intrinsic optical signal imaging of hemodynamic signals from the cortex during locomotion. A) Location of the frontal, forelimb/hindlimb whisker and visual cortex regions of interest (ROI). All but the frontal area are determined from cytochrome oxidase staining. Scale bar denotes 1mm. B) Changes in reflectance obtained at 530 nm over the entire cranial window at different time points during the imaging session. Decreases in reflectance correspond to increases in blood volume, caused by vessel dilation. C) Top, change in reflectance in each of the four ROIs versus time. The black ticks show locomotion events. There is a large decrease in reflectance, corresponding to vasodilation, in all ROIs but the frontal cortex during locomotion. Note the spontaneous fluctuations in blood volume at rest. Middle, spectrogram of the intrinsic optical signal over the entire hemisphere showing heart-rate fluctuations. The heart-rate is visible as a peak in the spectrogram, and increases during the locomotion bout. The blue line shows the peak of the heart-rate. Bottom, spectrogram of the electrocortigram (ECoG) versus time. Locomotion is accompanied by increases in high frequency (gamma-band) power. Gamma band power, changes versus resting, in dB, are plotted in green. D) Locomotion-triggered average from the entire 30 minute trial. Note the strong responses in the FL/HL, whisker and visual cortex, but lack of response in frontal cortex. E) The arterial and venous components of the blood volume response to locomotion in each pixel can be separated using a linear fitting procedure into an arterial component and a venous component (Huo et al., 2015a). The amplitude of the arterial component for each pixel is plotted at the top (in red), and the venous amplitude (in blue) for each pixel is plotted versus the bottom. The brighter the color, the larger the amplitude of the fit to the pixel. The venous dilation is widely spread across the cortex, while the arterial response is more localized. Scale bar is 1mm.

Technical challenges involved in imaging awake animals in MR scanners include controlling for motion artifacts and minimizing the stress induced by scanning noise and environment. To resolve these issues, an acclimation procedure is typically applied to habituate the animal to MRI scanning environment. Through this acclimation paradigm, animal’s motion and stress during MRI scanning are substantially diminished (Ferenczi et al., 2016) (King et al., 2005). This method has been extensively used in a number of awake animal fMRI studies involving multiple species including mice (Desai et al., 2011; Harris et al., 2015), rats (Becerra et al., 2011; Ferris et al., 2006; Huang et al., 2011; Lahti et al., 1998; Liang et al., 2011; Martin et al., 2006; Zhang et al., 2010), monkeys (Belcher et al., 2013a; Hung et al., 2015b; Liu et al., 2013b), rabbits (Schroeder et al., 2016) and dogs (Andics et al., 2016; Berns et al., 2012).

Although acclimation paradigms vary across labs, they typically use a noninvasive restraint system and a routine habituation procedure over several days (see review (Febo, 2011)). A rat head restrainer is usually composed to a bite bar, a pair of adjustable ear bars or ear pads, a nose clamp and a shoulder pad. Such a device is commercially available (e.g. Animal Imaging Research, LLC, MA, USA). In terms of acclimation procedure, using our lab as an example (**Figure 11**), the rat is first briefly anesthetized (2-3% isoflurane given for 3-5 minutes) to enable secure placement of the animal’s head into a head holder with the canines secured over a bite bar, the nose fixed with a clamp, and ears positioned inside the head holder with adjustable ear pads or ear bars. Topical application of lidocaine (2%) or EMLA cream is used to relieve any discomfort associated with head-fixation. The animal’s forepaws and hindpaws are loosely taped to prevent any self-injurious behavior (i.e. scratching). The body is then placed in a Plexiglas body tube that allows unrestricted respiration and movement of the trunk and limbs. The entire unit is subsequently secured onto a firm base, and then placed in a black opaque tube “mock scanner”. An audio-recording of acoustic sounds from various imaging pulse sequences are played throughout the acclimation procedure. The animal is exposed to these conditions for 7 days before imaging. The time for exposure during acclimation is increased from 15 minutes on the first day to 60 minutes on days 4, 5, 6, and 7, with an increment of 15 minutes per day.

**Figure 11.**
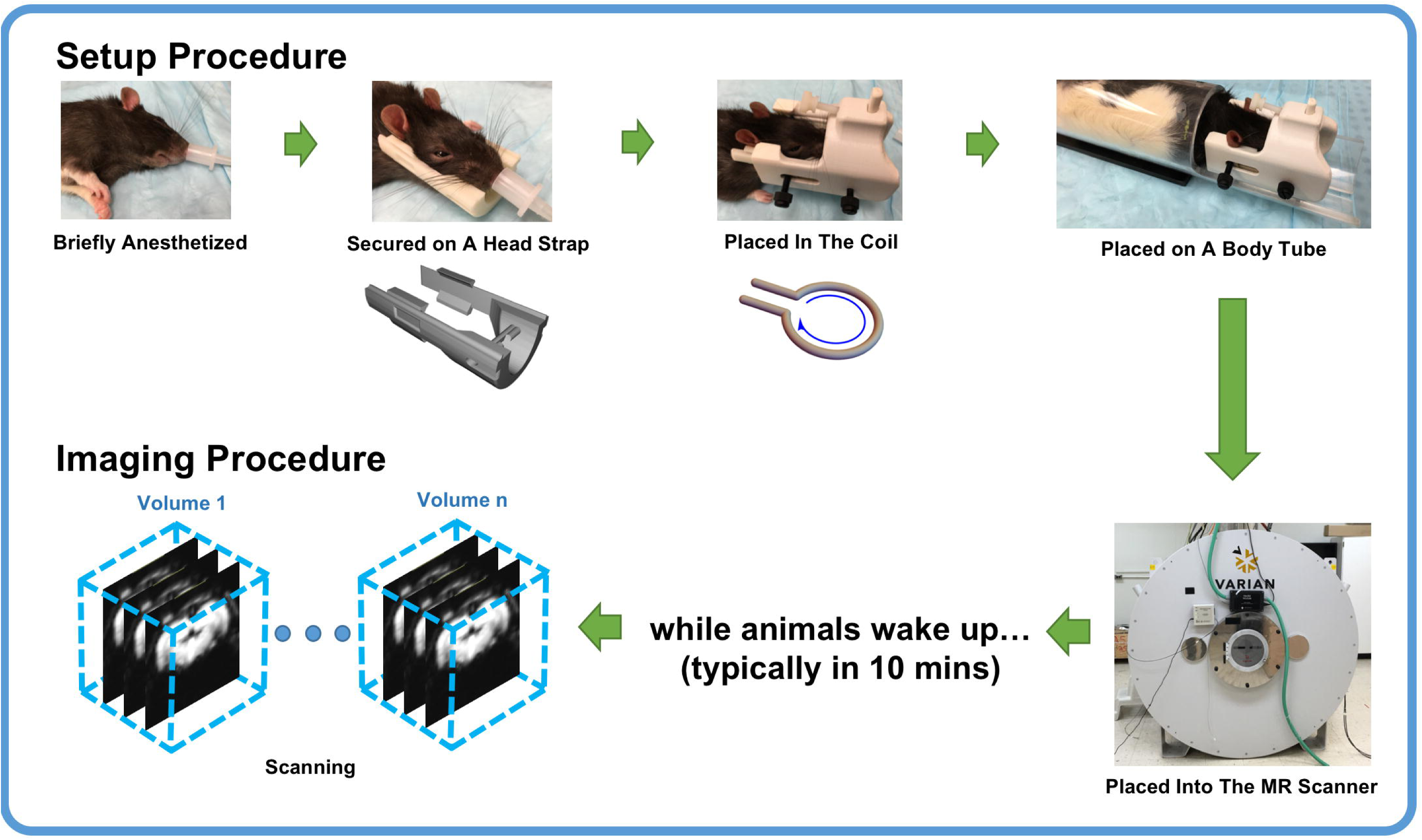
Procedure for imaging an awake rat using MRI. Schematic illustration of the animal setup and imaging procedures.

### Minimizing stress during awake imaging

One concern with the use of awake animals is that they might be overly stressed by head-fixation or imaging. While we cannot say that there is no stress associated with awake imaging procedures, this stress is most likely minimal. For optical imaging in mice, habituation for several days is likely on the conservative side, as some groups successfully use as short as one day of acclimation (Sofroniew et al., 2014), animals become proficient in using a treadmill in 10-15 minutes (Niell and Stryker, 2010), and habituation does not change neural responses, as neural activity in the motor cortex is the same in habituated and un-habituated mice (Dombeck et al., 2009). Mice rapidly (usually within one session) acclimate to head-fixation, and learn to ‘operate’ the treadmill. After acclimation, resting heart rates in head-fixed mice are comparable to those from recorded telemetrically from mice in their home cage (Gehrmann et al., 2000, Huo et al., 2015b) (**Figure 10**), indicating that that head fixation after acclimation is not stressful. In our hands, the pattern of evoked hemodynamic signals are stable up to 10 months of continuous imaging (Huo et al., 2015a, 2015b), which would not be the case if the animals were stressed. Many other groups have used head-fixed rodents, and have shown that they can perform complex, cognitively demanding tasks (Harvey et al., 2009; Komiyama et al., 2010; Mayrhofer et al., 2013; Mehta, 2007; Smear et al., 2011) that use memory and discrimination as stimuli at very high levels, which would not be possible if the animals were excessively stressed.

In awake animals, invasive measurements of physiological variables (arterial pressure, blood gasses, etc.) are more difficult to measure. However, other physiological parameters, such as heart rate (**Figure 10**), can be easily extracted from optically signals (Drew et al., 2011; Huo et al., 2015a, 2015b). Other indicators of state, such as whisker or body motion (**Figures 8**, **9** and **10**) can also be used as moment-to-moment measures of animal’s state.

Likewise, although restraint is involved in the MRI acclimation paradigm, this procedure is not likely to induce chronic stress in animals. Unlike studies investigating the chronic stress effects of immobilization, the restraint for acclimation and imaging requires considerably shorter restraining period per day and only involves restraint of the head as oppose to the whole body. Indeed, King et al. reported that the level of stress hormone (corticosterone) and other physiologic parameters of rats return to pre-stress baseline following 5-8 days of acclimation (King et al., 2005). The data are consistent with reports that heart rate and blood pressure normalized after initial exposure to restraint in rats, indicating that the animal well adapted to the restrainer (Parry and McElligott, 1993). Rats that were acclimated and imaged up to 4 times did not show significantly altered anxiety level compared to intact rats as measured by elevated plus maze test (**Figure 12**).

**Figure 12.**
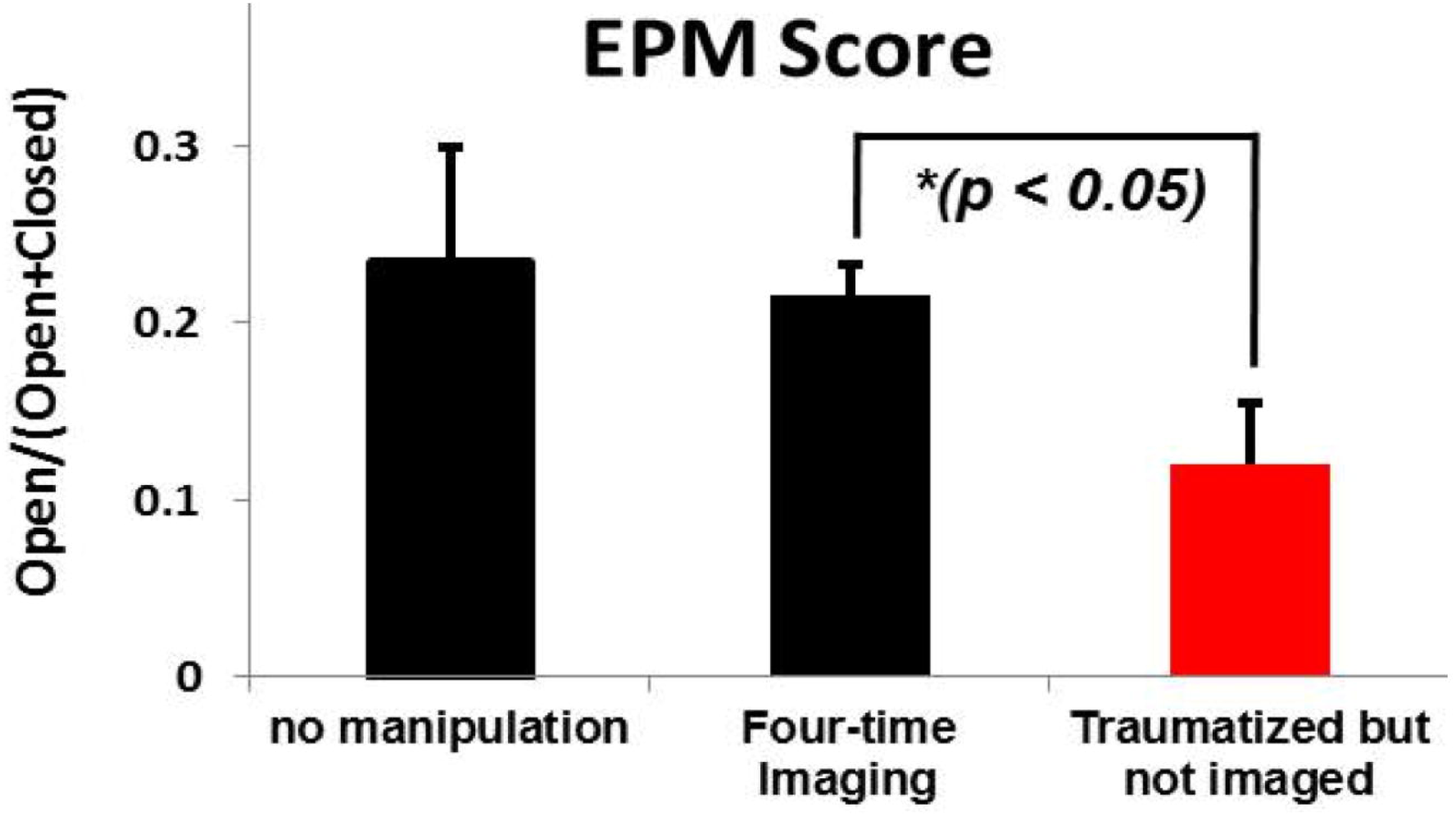
Stress induced by imaging is minimal. The elevated-plus maze (EPM) score (open/open+closed arm) was separately evaluated in three groups of rats: rats without any manipulation (n=8), rats imaged four times and rats exposed to a single episode of predator odor (i.e. traumatized) but did not undergo any acclimation and imaging procedures (n=8). EPM in the last two groups was measured 7 days after either the trauma exposure or the last imaging. Two sample t-tests showed that there was no significant difference between the no manipulation group and four-time imaging group (p=0.8), while the predator odor exposed group showed a significantly higher stress level relative to the imaging group (p=0.03). These results suggest that predator odor significantly increased the stress level while multiple imaging acquisitions had no significant effect. Bars show SEM.

Importantly, acclimation does not compound effects of other stressors on animals. In our recent study, the stress attributed to acclimation was minimal and significantly less than the stress introduced by a different stressor (i.e. single episode of predator odor exposure) (Liang et al., 2014). Notably, we found that there was no interaction between the acclimation-related stress and predator odor-related stress, suggesting that any possible residual effects from acclimation-and/or imaging-related stress can be subtracted out in testing animals with appropriate controls, which undergo the identical acclimation and/or imaging procedures. These results allow us to specifically examine the effect predator odor stress on neural circuit function in awake rodents (Liang et al., 2014). Taken together, these data suggest that stress induced by acclimation and imaging is very mild in animals and can be well controlled.

It is important to keep in mind that anesthesia is not ‘stress-free’, and in some cases can induce stress-related hormonal changes substantially larger than behavioral stressors. Urethane increases circulating adrenocorticotropic hormone by a factor of four, substantially higher than open-field stress (Donnerer and Lembeck, 1988). Studies in human patients have shown that anesthesia increases circulating cytokynes (Brand et al., 1997), and can increase circulating epinephrine, norepinephrine, and cortisol (Kono et al., 1981) by up to a factor of two or more, though fentanyl can cause a substantial decrease in circulating cortisol relative to control conditions (Kono et al., 1981). Above and beyond the effects of anesthetic, surgical procedures can cause enormous hormonal changes (Desborough, 2000), making it preferable to temporally separate any surgical procedure (such as cranial window implantation) from the imaging session.

### Neurovascular coupling and functional connectivity are robust and stable in the awake animal

Historically, anesthesia has been used as an important experimental tool for perturbing the state of the brain in a reproducible manner. For awake animal imaging approaches to be successfully applied, it is essential to validate the reproducibility of data collected.

Neuroimaging data collected in awake animals using aforementioned techniques demonstrate a high degree of reproducibility. Our opto-fMRI data acquired in two sessions separated by 2-7 days show highly consistent BOLD activation patterns in all rats (**Figure 13a**). Besides, BOLD amplitude of the corresponding voxels in the two maps were highly correlated (r = 0.79, p<10^-20^) (Liang et al., 2015b). Similarly, rsfMRI data collected in two subgroups of awake rodents show highly reproducible connectivity profile, as well as a high correlation in voxel-wise connectivity strength (**Figure 13b**) (Liang et al., 2012a). Similarly, when locomotion-induced changes in CBV are fit with linear models, these fits are very reproducible across days (Huo et al., 2015a) (**Figure 13c,d**), indicating that even with spontaneous, volitional behaviors, the hemodynamic response is stable in head-fixed animals.

**Figure 13.**
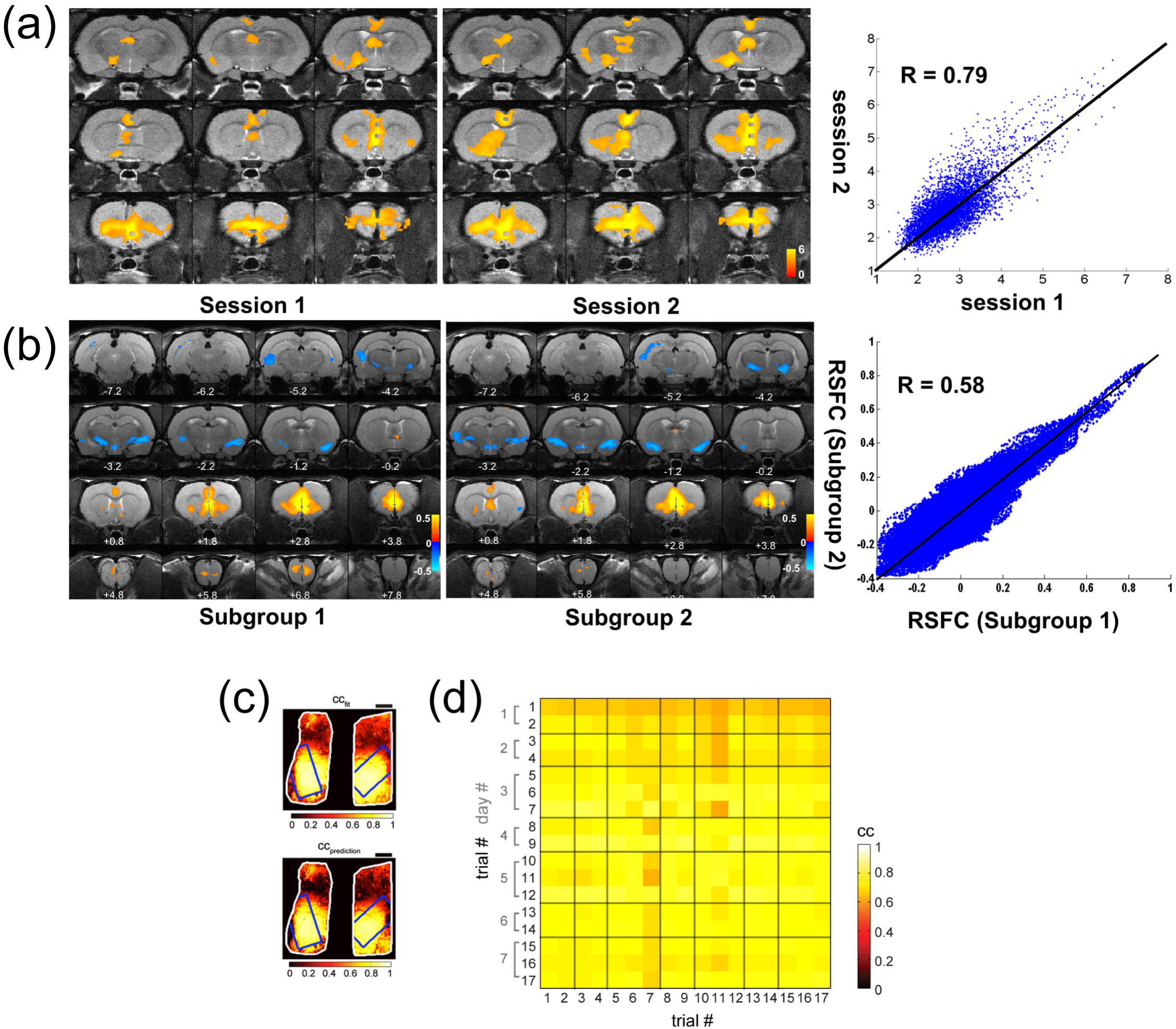
Functional networks and hemodynamic responses are reproducible and stable over time. A) bold activation maps in response to optogenetic stimulation of IL in awake rats on two separate days. Left panel: bold activation map acquired on one day; middle panel: bold activation map acquired in the same group of awake animals on another day; right panel: correlation of voxel-wise bold amplitude between the two days. Adapted from (Liang et al. 2015). B) IL RSFC maps in two separate subgroups of animals. Left panel: IL RSFC map in subgroup 1; middle panel: IL RSFC map in subgroup 2; right panel: correlation of voxel-wise RSFC strength between the two subgroups. Adapted from (Liang et al. 2011). C) Top, the correlation coefficient for each pixel in the window between the predicted hemodynamic response to locomotion (using a fitted hemodynamic response function) during one 30-minute IOS imaging session. The blue rectangles denote histologically-identified forelimb/hindlimb representations. The fit has a high correlation in these regions, and not in the frontal cortex because there is no hemodynamic response in the frontal cortex to locomotion. Bottom shows the correlation between the predicted hemodynamic response and actual response plotted for each pixel for a different 30-minute imaging session, using the fits same in the top panel, demonstrating a high reproducibility of the response. D) Matrix of cross-correlations between the predicted and actual hemodynamic response to voluntary locomotion during a 30-minute trial for the histologically defined SI FL/HL area. Fits from the trials numbered in the rows were used to predicted data from the trials in the columns. Imaging sessions were done on seven different days with 17 days between the first and last day, showing that the hemodynamic responses are stable. C and D adapted from (Huo et al. 2015a).

One factor that may confound the reproducibility of neuroimaging data in awake animals is spontaneous behaviors. Any awake animal (or human) will engage in small spontaneous behaviors in the scanner, and these spontaneous behaviors could affect hemodynamic signals. Several behaviors known to cause or correlate with measurable changes in hemodynamic signals and functional connectivity include eye state (closed, open or fixating) (Patriat et al. 2013; Chang et al., 2016), blink rate (Guipponi et al., 2014), and variations in respiration (Birn et al., 2009). In rodents, behaviors like whisking and sniffing (Moore et al., 2013) are analogous to eye movements in humans and primates (Kleinfeld et al., 2006; Crapse and Sommer, 2008; Ahissar and Assa, 2016), and these behaviors would be expected to have similar impacts on functional connectivity. These behavioral effects can be regressed out, as is currently done with physiological noise (Birn 2012; Murphy et al., 2013). Going forward, monitoring of the relevant behavior(s) (Figure 8 and (Guipponi et al., 2014; Chang et al., 2016; McGinley et al., 2015; O’Connor et al, 2010; Yang et al., 2016)) in animals is likely to play an important role in animal fMRI experiments, just as it has in human experiments (Richlan et al., 2014; Wang et al., 2016). The development of new techniques for analyzing behavioral data in other paradigms have undergone enormous advances in recent years (Gomez-Marin et al. 2014), and it is likely that some of these approaches can be applied to the studies of more subtle behaviors.

Another concern is the normal fluctuations in blood pressure and heart rate that are found in the awake state (Goldberger et al., 2002) might affect neurovascular coupling. However, sensory-evoked changes in CBF and arterial dilation are not affected by large changes in heart rate in the awake animal. CBF measurements in mice where heart rate was pharmacologically elevated with glycopyrrolate, or decreased with atenolol showed that sensory-evoked CBF increase was not affected by these large manipulations of heart rate, and neither was arterial dilation (Huo et al., 2015b). However, pharmacological depression of heart rate reduced venous dilation in response to sensory stimulation, similar to that in the anesthetized animal (Drew et al., 2011). These results indicate that in the awake animal, the hemodynamic response is largely un-affected by cardiovascular fluctuations as long as the heart rate is not depressed. Further work is still needed to understand what contributions normal cardiovascular fluctuations might have on neurovascular coupling.

## Conclusions

Anesthetics have been widely used for studying neurovascular coupling in *in-vivo* animal models, and these models have added tremendously to our understanding of neurovascular coupling and associated physiological processes (Logothetis et al., 2001; Vincent et al., 2007; Shulman et al., 1999). However, anesthesia has profound actions on vascular, electrical, metabolic, and other physiological processes, and alters the coupling relationship between neural and vascular responses. Therefore, it needs to be cautious when one aims to translate functional neuroimaging data acquired in anesthetized animals to awake humans. Recent technical developments have enabled the use of optical and MR imaging in awake and behaving animals, allowing scientists to address critical questions about neurovascular coupling and brain-wide circuit function in a preparation similar to the awake human. Therefore, we believe the time is right for the community to move towards using awake animal paradigms when addressing questions involving neurovascular coupling and circuit function.

## Conflict of interest

none

## Acknowledgments

PJD is supported by an award from the American Heart Association, a Scholar Award from the McKnight Endowment Fund for Neuroscience, and R01NS078168, R01EB021703, and R01NS079737 from the NIH. NZ is supported by R01MH098003 from the National Institute of Mental Health and R01NS085200 from the National Institute of Neurological Disorders and Stroke.

